# Metabolic dysfunction related to specific genetic variant sites of methylmalonic acidemia

**DOI:** 10.64898/2025.12.15.694311

**Authors:** Haijuan Zhi, Yuxin Deng, Yi Ding, Ting Chen, Bohui Zhou, Yue Cao, Siyu Chang, Xia Zhan, Feng Xu, Lili Hao, Lili Liang, Wenjuan Qiu, Huiwen Zhang, Xuefan Gu, Lianshu Han

## Abstract

Methylmalonic acidemia (MMA) is a genetic metabolic disease, with mut (mut-MMA, caused by *MMUT* gene variation) and cblC (cblC-MMA, caused by *MMACHC* gene variation) being the predominant subtypes. MMA patients carrying c.729_730insTT and c.609G>A exhibit severe clinical manifestations and have poorer outcomes. Little information is known on the potential mechanism. The metabotypes of MMA patients carrying c.729_730insTT and c.609G>A were profiled. The potential effect of genetic variation sites on the metabolism were explored. Leveraging the gene variation spectrum and the targeted metabolomics, propionic acid, indolelactic acid, myristoleic acid, tetradecenoic acid, tridecanoic acid, and oxoadipic acid showed strong enrichment in c.729_730insTT carrying mut-MMA patients. Whereas mutation c.609G>A caused adipic acid, pimelic acid, fructose, AMP, hexanylcarnitine, and methionine significnatly altered in cblC-MMA patients. Furthermore, the metabolomic profiles were obviously different in c.729_730insTT and non-c.729_730insTT carrying mut-MMA patients after treatment. Different metabolic alterations in cblC-MMA patients carrying c.609G > A and non-c.609G > A under treatment were also observed. Our findings provide unique insights into the metabolic basis of MMA specific subtypes and its implications across different variants.

## Introduction

Methylmalonic acidemias (MMA) comprises a series of organic acid metabolic disorders that are mainly inherited in autosomal recessive mode.^1–3^ The global average incidence of MMA is about 1:100,000 and ranges from 1:3500 to 1:39,000 in China.^4–6^ MMA is caused by disorders of methylmalonyl-CoA mutase or cobalamin metabolism and is characterized by the accumulation of methylmalonic acid and other toxic metabolites in the body, which could cause various clinical symptoms through complex mechanisms.^3,7,8^ The clinical phenotypes of MMA are variable and atypical, with onset ranging from prenatal to adulthood and clinical symptoms ranging from minor to life-threatening.^7,9–12^

MMA can be divided into combined and isolated types according to whether homocysteinemia is combined biochemically.^7^ In China, 60%–80% of MMA cases are combined, with the cblC subtype (cblC-MMA, caused by *MMACHC* gene mutations) predominant (95%) and mut subtype (mut-MMA, caused by *MMUT* gene mutations) account for 90% of the isolated MMA population.^10,13^ The *MMUT* gene carries a wide variety of mutations. c.729_730insTT was the most frequent *MMUT* mutation in China and the presence of c.729_730insTT was associated with more poor outcomes.^10,13^ Our previous study pointed out that c.729_730insTT-carrying patients exhibited a low treatment effective rate and a poor prognosis.^13–15^ Additionally, the mutation c.729_730insTT was found in patients not responsive to vitamin B12.^16^ And c.609G>A was the most frequent *MMACHC* mutation detected in Chinese patients and was independently associated with poor outcomes, especially for neurodevelopmental deterioration.^17,18^ Furthermore, the variant of c.609G>A was increasingly observed in symptomatic patients.^19^ At present, there is no information on the potential underlying mechanism explaining why the mutation c.729_730insTT and c.609G>A causes the poorer phenotype.

Metabolite represents a molecular readout of an underlying (patho-)physiological process.^20–23^ Human metabolites play a crucial role in maintaining physiological homeostasis and health.^24,25^ Circulating metabolite levels are strongly influenced by dietary habits, lifestyle, and genetic background.^26,27^ Many metabolites exhibit high heritability and are proximal to the clinical endpoints, making them effective intermediate phenotypes that link genetic susceptibility and diseases.^27^ Revealing metabolic underpinnings under different genetic background helps provide insights into the disease etiology. Metabolic reprogramming emerges as a critical mechanistic link that orchestrates genotypic and metabotypic variabilities.^28^ Additionally, elucidating metabolic alterations is essential for identifying early intervention targets and developing accurately preventive strategies for MMA with specific mutation sites. ^22,29,30^ Metabolomics providing a direct perspective on biological metabolism could reflect the functional consequences of gene expression changes, and contributing to a more comprehensive understanding of disease mechanisms.^31,32^ A range of high-throughput technologies now enable examination of the genetic regulation of biochemical individuality at the population scale.^25,33,34^ Existing untargeted and targeted platforms provide highly synergistic information due to limited overlap in the metabolome coverage.^35,36^ Liquid chromatography-mass spectrometry (LC-MS) metabolomics approaches measure a wide range of metabolites, including amino acids, organic acids, nucleotides, lipids, and other low-molecular-weight metabolites that cover comprehensive biological functions and pathological pathways.^35–37^ In addition, several metabolomic studies on MMA have conducted based on small sample size and a few detected metabolites.^4,38–43^ Nevertheless, it remains challenging to ascertain the origin(s) of metabolic changes and elucidate gene mutations specifically contribute to metabolic alterations of MMA. Furthermore, MMA patients with specific mutations (c.729_730insTT and c.609G>A) present different clinical symptoms and prognosis, and the exact influencing factors and mechanisms are still unclear.

This study aims to summarize the relationships between specific MMA gene mutation sites and metabotypes. In this retrospective study, based on the genetic records of a large MMA cohort, we analyzed metabolic profiles and the influence of specific mutation sites on the metabolic features of mut-MMA and cblC-MMA patients. Herein, we performed a targeted metabolic analysis to determine the plasma metabolites and the biological pathways associated with MMA gene mutations (c.729_730insTT or c.609G > A mutations). Furthermore, we assessed the potential role of treatment on the metabolomic signature of the patients with specific mutations (c.729_730insTT or c.609G > A). Our findings will allow us to gain a better understanding of the mechanisms through which specific gene mutation carrying leads to poorer outcomes in MMA and enhance the understanding of metabolic architecture underlying MMA subtypes across diverse genetic backgrounds.

## Methods

### Study population

A total of 117 MMA patients underwent genetic analysis were enrolled from January 2016 to December 2024. Among them, 59 cases (before and after treatment, n = 29; only before treatment, n = 11; only after treatment, n = 19) were caused by the *MMUT* gene variations and 58 cases (before and after treatment, n = 19; only before treatment, n = 20; only after treatment, n = 19) were caused by the *MMACHC* gene variations (Supplementary Tables 1-7, Supplementary Figure 1). All of these patients were followed up at least once and included in our study. This study was in accordance with the Helsinki Declaration and has been approved by the Ethics Committee of Xinhua Hospital, Shanghai Jiaotong University School of Medicine (Approval ID: XHEC-D-2025-090).

Dried blood spots (DBSs) were collected and the levels of propionylcarnitine (C3), acetylcarnitine (C2), and methionine (Met) were analyzed using tandem mass spectrometry (API4500, Applied Biosystems, Foster City, CA).^9^ Genomic DNA was extracted from peripheral blood (Qiagen Blood DNA Mini Kits, Hilden, Germany). Sanger sequencing or next generation sequencing were performed and variations were identified using reference sequences from Genbank (*MMUT*: NC_000006.12; *MMACHC*: NM_015506). For the variants that were not recorded in the Human Gene Mutation Database, the ClinVar Database (https://www.ncbi.nlm.nih.gov/clinvar/) or previous literatures, the pathogenicity analysis of these variants was performed according to the American College of Medical Genetics and Genomics and the Association for Molecular Pathology (ACMG-AMP) guidelines.^7,13^

### DBS metabolomic analysis

Targeted metabolomic profiling based on liquid chromatography coupled with tandem mass spectrometry (LC-MS/MS) for DBS was performed utilizing a Q300 kit (Metabo-Profile, Shanghai, China). Briefly, each DBS sample was mixed with 150 μL ice-cold methanol with internal standards added, 20 μL of deionized water, and vortexed for 5 min, followed by centrifuged (4000 g for 30 min at 4 ℃). Then 30 μL supernatant was transferred to a 96-well plate and 20 μL of freshly prepared derivative reagent was added to each well (30 ℃ for 60 min). Afterwards, 330 μL of 50 % ice-cold methanol solution was added (incubated, −20 ℃ for 20 min). Then the plated was centrifuged at 4000 g at 4 ℃ for 30 min. And 135 μL of supernatant was transferred to a new 96-well plate with 10 μL internal standards added in each well and used for subsequent LC-MS/MS analysis.^44,45^

### Statistical analysis

The processed metabolomic data were subjected to multivariate analysis after normalization, including scaled principal component analysis (PCA) and orthogonal partial least-squares discriminant analysis (OPLS-DA) analysis, utilized to identify metabolomic alterations. The cross-validation and response permutation tests were performed to evaluate the robustness of the model. The variable importance in projection (VIP) value of each variable in the OPLS-DA model was calculated to indicate its contribution to classification. Significance was determined using an unpaired Student’s t-test. VIP values > 1, fold change (FC) > 1 and *p* < 0.05 were considered to indicate statistical significance. A heatmap analysis was conducted to visualize the differential expression of metabolites in mut (carrying c.729_730insTT or non-c.729_730insTT) and cblC subtypes (carrying c.609G > A or non-c.609G > A). Statistical and pathway analysis were performed using the online software MetaboAnalyst 6.0 (https://www.metaboanalyst.ca). Correlation analysis was performed using the Metware Cloud, a free online platform for data analysis (https://cloud.metware.cn).

## Results

We included 117 MMA patients from Xinhua Hospital Affiliated to Shanghai Jiao Tong University School of Medicine. Metabolomic profiling was performed using LC-MS/MS. PCA was performed to assess the quality of the data set. After quality control, 205 directly quantifiable metabolites and reaction ratios, primarily amino acids (18.54%), fatty acids (18.05%), organic acids (12.20), and related reaction ratios (12.20%) were included in the subsequent analyses (Supplementary Table 8, Supplementary Figure 2A, B).

### Overview of metabolomic results for MMA with different genetic background

Metabolomic analysis of mut and cblC MMA subtypes was comprehensively performed. Firstly, DBS metabolites with significant differences between mut and control were defined using the multi- and univariate analysis (Supplementary Figure 3A-D, Supplementary Table 9). In addition, spearman rank correlation analyses were used to evaluate the relationship between the significantly altered metabolites (Supplementary Figure 3E, Supplementary Table 10). Propionylcarnitine, methylmalonylcarnitine, and methylmalonic acid were negatively correlated with glutamic acid, glyceric acid, N-acetylglucosamine, arachidonic acid, 4-hydroxybenzoic acid, fructose, phenylpyruvic acid, and phenylpyruvic acid/ortho-hydroxyphenylacetic acid. Propionylcarnitine, methylmalonylcarnitine, and methylmalonic acid were positively correlated with valerylcarnitine, hydroxypropionic acid, 3-hydroxyisovaleric acid, 2-methylbutyroylcarnitine, acetylcarnitine, asparagine/aspartic acid, 2-furoic acid, lactic acid/pyruvic acid, butyrylcarnitine, hexanylcarnitine. Pathway analysis of the 66 selected potential biomarkers showed that significantly enriched pathway mainly involved glyoxylate and dicarboxylate metabolism, alanine, aspartate and glutamate metabolism, arginine biosynthesis, pentose phosphate pathway, and glycine, serine, and threonine metabolism (Supplementary Figure 3F, Supplementary Table 11).

Next, DBS metabolites with significant differences between cblC and control were obtained based on the OPLS-DA model and volcano plot (FC cutoffs >1 or <1, and significance *p*-values < 0.05) (Supplementary Figure 4A-D, Supplementary Table 12). Subsequently, 60 significantly altered metabolites were subjected to correlation analysis. Propionylcarnitine was negatively correlated to glutamic acid, oxoadipic acid/aminoadipic acid, 4-hydroxybenzoic acid, N-acetylglucosamine, indole-3-carboxaldehyde, phenylpyruvic acid, oxoadipic acid, indole-3-carboxylic acid, linoleic acid, arachidonic acid, and pyruvic acid. Methylmalonylcarnitine was negatively correlated to glutamic acid, oxoadipic acid/aminoadipic acid, 4-hydroxybenzoic acid, and phenylpyruvic acid. Methylmalonic acid was negatively correlated to glutamic acid, linoleic acid, arginine, and phenylpyruvic acid. Propionylcarnitine, methylmalonylcarnitine, and methylmalonic acid were positively correlated with lactic acid/pyruvic acid and N-alpha-acetyl-L-lysine (Supplementary Figure 4E, Supplementary Table 13). Functional analysis revealed that the altered pathways in cblC mainly involved in phenylalanine metabolism, phenylalanine, tyrosine and tryptophan biosynthesis, alanine, aspartate and glutamate metabolism, and arginine biosynthesis (Supplementary Figure 4F, Supplementary Table 14).

To further examine the metabolomic variance between cblC and mut, additional OPLS-DA model was built (Supplementary Figure 5A). Subsequently, a differential analysis of all detected metabolites or reaction ratios (FC cutoffs >1 or <1, and significance *p*-values <0.05) was performed (Supplementary Figure 5B). We defined 51 potential biomarkers based on VIP, FC, and *p* value (Supplementary Figure 5C, D, Supplementary Table 15) and identified significant associations between these biomarkers. Propionylcarnitine was negatively correlated with dimethylglycine and 2-methy-4-pentenoic acid. Methionine was negatively correlated with succinic acid/fumaric acid. Positive association between methylmalonylcarnitine and gluconic acid/gluconolactone was observed (Supplementary Figure 5E, Supplementary Table 16). Addiontionally, for cblC vs mut, one carbon pool by folate, glycine, serine and threonine metabolism, arginine biosynthesis, amino sugar and nucleotide sugar metabolism, and fructose and mannose metabolism were among the most affected pathways (Supplementary Figure 5F, Supplementary Table 17).

IEMs are metabolic diseases caused by rare genetic variants that lead to metabolite deficiency and/or accumulation, resulting in specific metabolic phenotypes. To explore the potential pathogenesis of MMA associated with specific genetic mutations (c.729_730insTT or c.609G > A), subsequently, we performed comparative metabolomics by leveraging the genetic mutation spectrum (*MMUT* and *MMACHC*, Supplementary Figure 1) and the quantified metabolites based on the dimension-reduction statistical analysis.

### Metabolomic profiles of c.729_730insTT and non-c.729_730insTT carriers of mut-MMA

To identify specific gene variant-associated metabotypes in mut-MMA patients, we applied OPLS-DA model as dimension-reduction approach to extract metabolomic signatures of c.729_730insTT and non-c.729_730insTT carriers before and after treatment, respectively.

Among the detected metabolites or reaction ratios, 66 were selected as the potential biomarkers between c.729_730insTT and non-c.729_730insTT carriers of mut-MMA before treatment based on the FC, *p* value, and VIP (Fig. 1A-D, Supplementary Table 18). The significantly up-regulated in c.729_730insTT carriers were assigned to propionic acid, indolelactic acid, pipecolic acid, 9E-tetradecenoic acid, etc., and down-regulated assigned to indole-3-propionic acid, phenylalanylphenylalanine, 2-hydroxyglutaric acid/oxoglutaric acid, etc. (Supplementary Table 18). Propionylcarnitine was negatively correlated with glycolic acid, lactic acid/pyruvic acid, and 2-methy-4-pentenoic acid (Fig. 1E, Supplementary Table 19). Positive associations between different classes of metabolites were observed, indicating synergistic effects between these processes. To further investigate the potential effect of c.729_730insTT on metabolite alterations, we conducted KEGG analysis. Valine, leucine and isoleucine biosynthesis and degradation pathways were the most affected. Alanine, aspartate and glutamate metabolism and lysine degradation were also significantly altered (Fig. 1F, Supplementary Table 20). Investigation of metabolomic differentiation between c.729_730insTT and non-c.729_730insTT carriers after treatment were also performed. Differential analysis based on the OPLS-DA and volcano plot of all metabolomic signatures was performed, resulting in 14 potential biomarkers (Fig. 2A-D, Supplementary Table 21). The main significantly altered biomarkers were mainly fatty acids, amino acids, and organic acids, including myristoleic acid, N-acetytyrosine, oxoadipic acid, etc. Oxoadipic acid was positively correlated with pyruvic acid (Fig. 2E, Supplementary Table 22). Pathway analysis of the 14 potential biomarkers showed that significantly enriched pathway mainly involved pyruvate metabolism, lipoic acid metabolism, glycolysis or gluconeogenesis, cysteine and methionine metabolism, and citrate cycle (Fig. 2F, Supplementary Table 23).

**Fig. 1:**
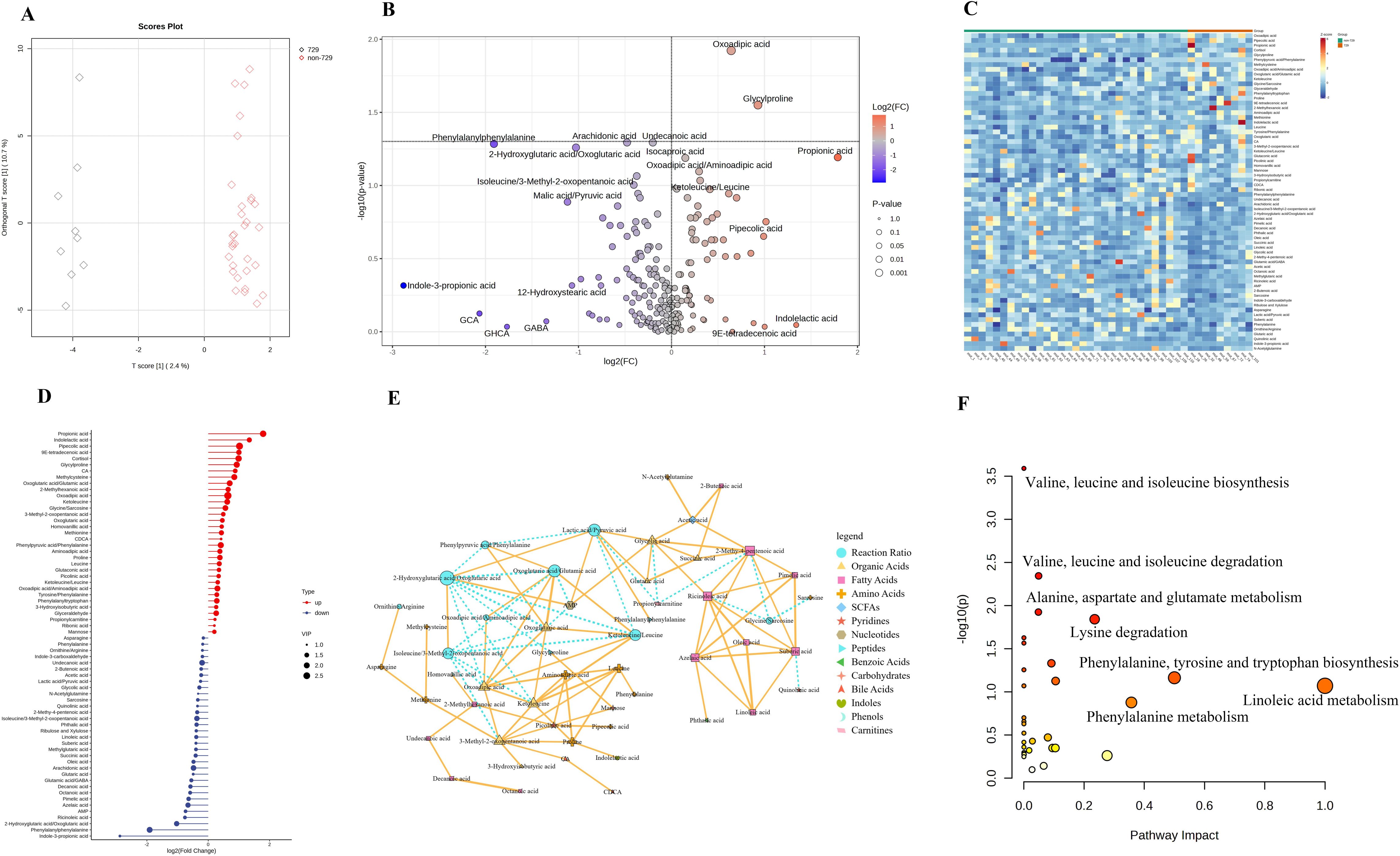
Comparative metabolome analysis of c.729_730insTT and non-c.729_730insTT carriers of mut. (A) OPLS-DA score plot. (B) Volcano plot showing the differential metabolites. The horizontal dashed line corresponds to the significance threshold of p = 0.05. (C) Heatmap analysis of the potential biomarkers. Potential biomarkers selected based on OPLS-DA model and univariate statistics. Red represents upregulated metabolites, and blue represents downregulated metabolites. (D) Stickman plot showing the FC and VIP values of the potential biomarkers. (E) Correlation analysis of the potential biomarkers with significant differences (*p* < 0.05). The solid lines represent negative correlation, while the dotted lines represent positive correlation. The size of the nodes represents the connectivity of the biomarker, and the larger the connectivity, the larger the node. The thickness of the lines represents the magnitude of the absolute value of the correlation, and the larger the absolute value of the correlation, the thicker the line. (F) Pathway analysis based on the potential biomarkers. The x-axis represents pathway impact generated from topological analysis; the y-axis represents the corresponding *p* values. Con, control; FC, fold change; mut, patients with methylmalonyl-CoA mutase defect, OPLS-DA, orthogonal partial least squares discriminant analysis; VIP, variable importance in projection.

**Fig. 2:**
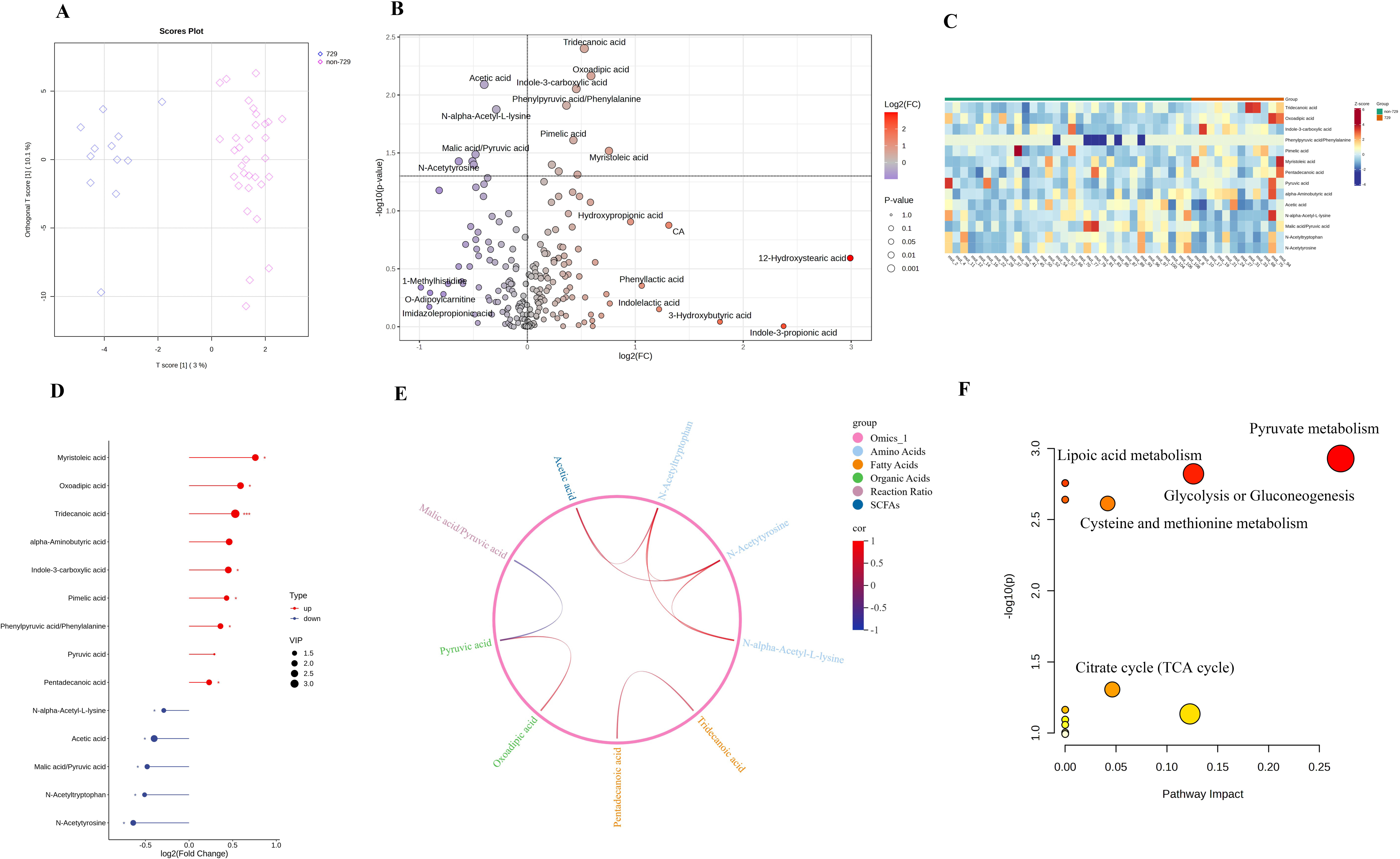
Comparative metabolome analysis of c.729_730insTT and non-c.729_730insTT carriers of mut after treatment. (A) OPLS-DA score plot. (B) Volcano plot showing the differential metabolites. The horizontal dashed line corresponds to the significance threshold of p = 0.05. (C) Heatmap analysis of the potential biomarkers. Potential biomarkers selected based on OPLS-DA model and univariate statistics. Red represents upregulated metabolites, and blue represents downregulated metabolites. (D) Stickman plot showing the FC and VIP values of the potential biomarkers. (E) Correlation analysis of the potential biomarkers with significant differences (*p* < 0.05). The Chord diagram was used to represent the correlation between various biomarkers. (F) Pathway analysis based on the potential biomarkers. The x-axis represents pathway impact generated from topological analysis; the y-axis represents the corresponding *p* values. Con, control; FC, fold change; mut, patients with methylmalonyl-CoA mutase defect, OPLS-DA, orthogonal partial least squares discriminant analysis; VIP, variable importance in projection.

Investigation of metabolomic shifts in response to treatment in c.729_730insTT and non-c.729_730insTT carriers were also performed, respectively. For mut-MMA patients carrying c.729_730insTT, OPLS-DA score plot shows that after treatment, the metabolomic profile was obviously altered (Fig. 3A). Based on the VIP value, *p* value, and FC, 29 potential biomarkers were classified (Fig. 3B-D, Supplementary Table 24). Fatty acids, incluidng pimelic acid, myristoleic acid, azelaic acid, and linoleic acid, were found to be significantly elevated in c.729_730insTT carriers after treatment, whereas oxoglutaric acid/glutamic acid, glycylproline, and oxoglutaric acid were significantly decreased (Supplementary Table 25). Correlation analyse among the potential biomarkers was performed (Fig. 3E, Supplementary Table 25). Significantly negative correlations were observed between indole-3-carboxaldehyde and ketoleucine/keucine, AMP and oxoglutaric acid/glutamic acid, linoleic acid and ketoleucine/leucine. Significantly positive correlations were observed between sarcosine and N-acetylglucosamine, imidazoleacetic acid and sarcosine, indole-3-carboxylic acid and suberic acid. KEGG analysis revealed that arginine biosynthesis, histidine metabolism, alanine, aspartate and glutamate metabolism, glycine, serine and threonine metabolism, and linoleic acid metabolism were the mainly altered pathway (Fig. 3F, Supplementary Table 26). For mut-MMA patients carrying non-c.729_730insTT, significant changes occurred in the metabolic profile after treatment (Fig. 4A-D). Significant features selected based on the VIP and volcano plot are listed in Supplementary Table S27. Carnitines, oleoylcarnitine, linoleylcarnitine, and carnitine, were significantly up-regulated in non-c.729_730insTT carriers after treatment. Bile acids (glycohyocholic acid, taurocholic acid, and cholic acid), fatty acids (adipic acid and 2-hydroxy-3-methylbutyric acid), organic acids (glutaric acid and 2-hydroxybutyric acid), and the ratio, oxoglutaric acid/glutamic acid were significantly down-regulated. Correlation analysis showed that ketoleucine/leucine was negatively associated with isoleucine/3-methyl-2-oxopentanoic acid and 2-hydroxyglutaric acid/oxoglutaric acid. Linolenic acid was negatively associated with 4-hydroxyproline/proline. Methylmalonylcarnitine was positively associated with linoleylcarnitine, oleoylcarnitine, and carnitine. Furthermore, chord diagram illustrated the correlations between different classes of metabolites, highlighting the internal correlations within carbohydrates and carnitines categories (Fig. 4E, Supplementary Table 28). The differential metabolites were mainly enriched in arginine biosynthesis, alanine, aspartate and glutamate metabolism, primary bile acid biosynthesis, and pentose phosphate pathway (Fig. 4F, Supplementary Table 29).

**Fig. 3:**
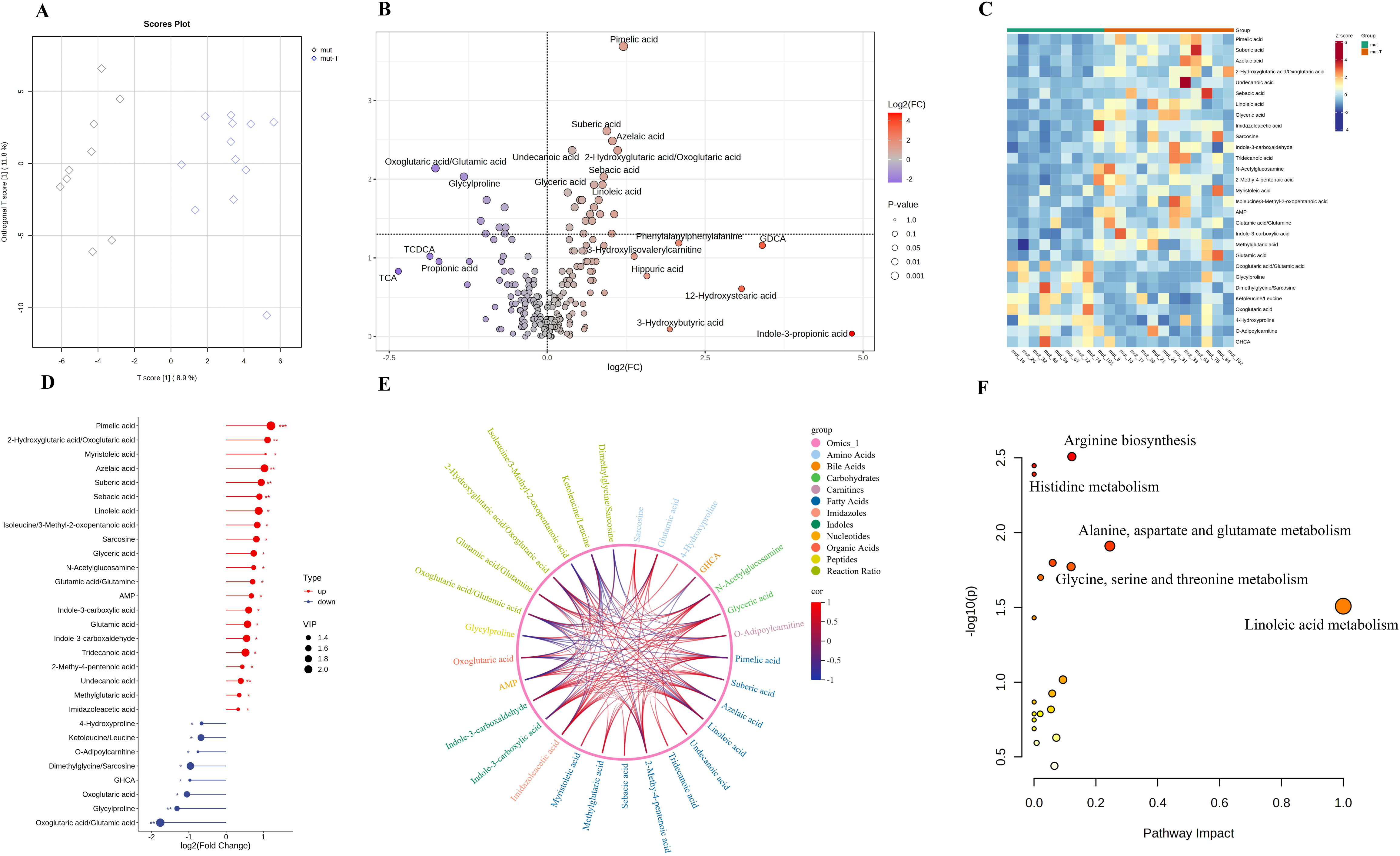
Metabolome alterations in c.729_730insTT carriers of mut after treatment. (A) OPLS-DA score plot. (B) Volcano plot showing the differential metabolites. The horizontal dashed line corresponds to the significance threshold of *p* = 0.05. (C) Heatmap analysis of the potential biomarkers. Potential biomarkers selected based on OPLS-DA model and univariate statistics. Red represents upregulated metabolites, and blue represents downregulated metabolites. (D) Stickman plot showing the FC and VIP values of the potential biomarkers. (E) Correlation analysis of the potential biomarkers with significant differences (*p* < 0.05). The Chord diagram was used to represent the correlation between various biomarkers. (F) Pathway analysis based on the potential biomarkers. The x-axis represents pathway impact generated from topological analysis; the y-axis represents the corresponding p values. Con, control; FC, fold change; mut, patients with methylmalonyl-CoA mutase defect, OPLS-DA, orthogonal partial least squares discriminant analysis; VIP, variable importance in projection.

**Fig. 4:**
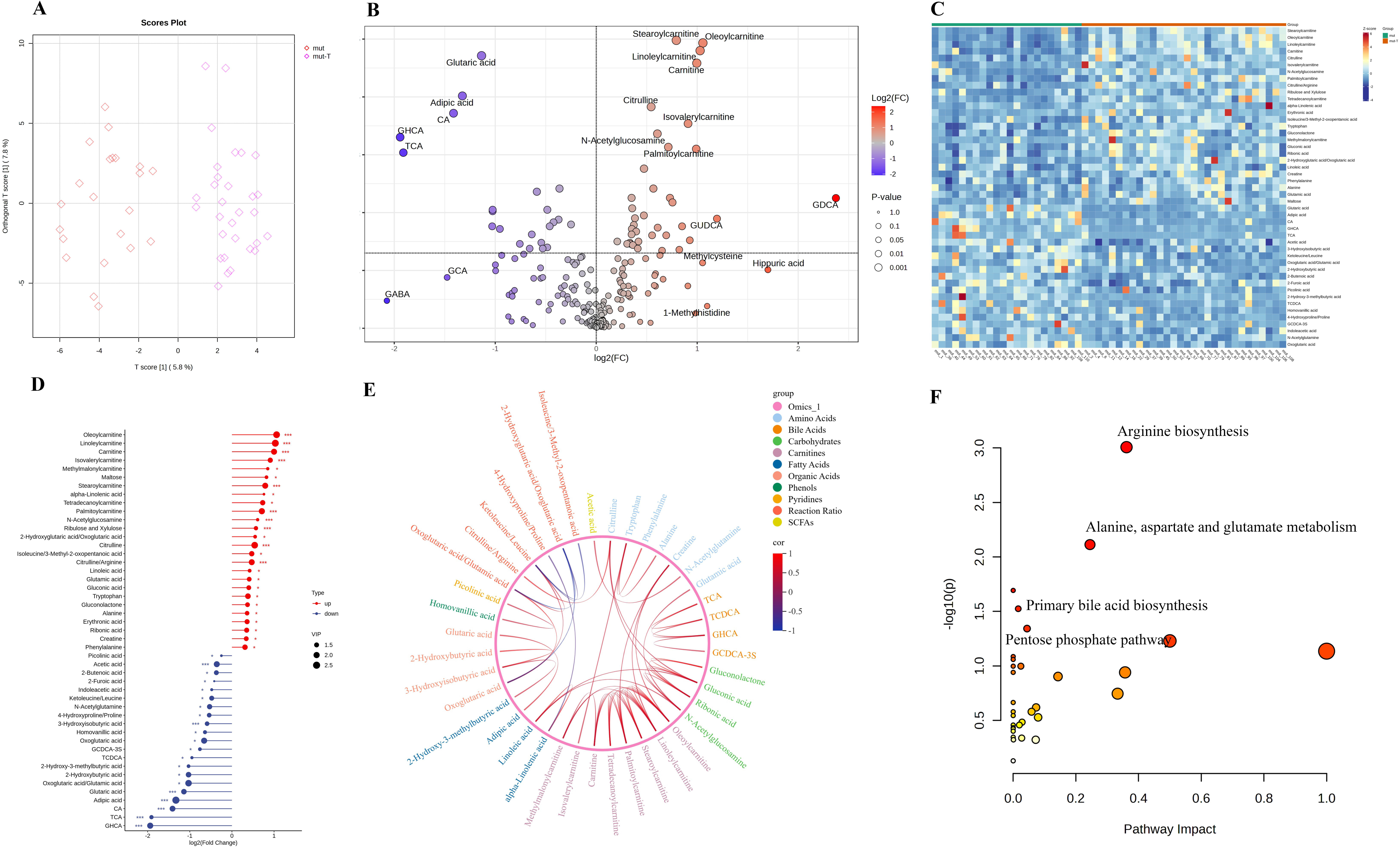
Metabolome alterations in non-c.729_730insTT carriers of mut after treatment. (A) OPLS-DA score plot. (B) Volcano plot showing the differential metabolites. The horizontal dashed line corresponds to the significance threshold of *p* = 0.05. (C) Heatmap analysis of the potential biomarkers. Potential biomarkers selected based on OPLS-DA model and univariate statistics. Red represents upregulated metabolites, and blue represents downregulated metabolites. (D) Stickman plot showing the FC and VIP values of the potential biomarkers. (E) Correlation analysis of the potential biomarkers with significant differences (*p* < 0.05). The Chord diagram was used to represent the correlation between various biomarkers. (F) Pathway analysis based on the potential biomarkers. The x-axis represents pathway impact generated from topological analysis; the y-axis represents the corresponding p values. Con, control; FC, fold change; mut, patients with methylmalonyl-CoA mutase defect, OPLS-DA, orthogonal partial least squares discriminant analysis; VIP, variable importance in projection.

The above results indicate that metabolomic phenotypes were significantly different between c.729_730insTT and non-c.729_730insTT carriers of mut-MMA whether before or after treatment. This imply that when treating mut-MMA patients, the specific gene mutation should be considered.

### Metabolomic profiles of c.609G > A and non-c.609G > A carriers of cblC-MMA

To investigate alterations in the metabolites of the cblC-MMA patients carrying c.609G > A and non-c.609G > A, metabolomics was also conducted. To examine the effect of gene mutation c.609G > A on the metabonotype, we generated OPLS-DA models with all 180 compounds and the calculated ratios extracted from the LC−MS/MS data set (Fig. 5-8A). DBS metabolomic signatures from different groups were obviously separated.

**Fig. 5:**
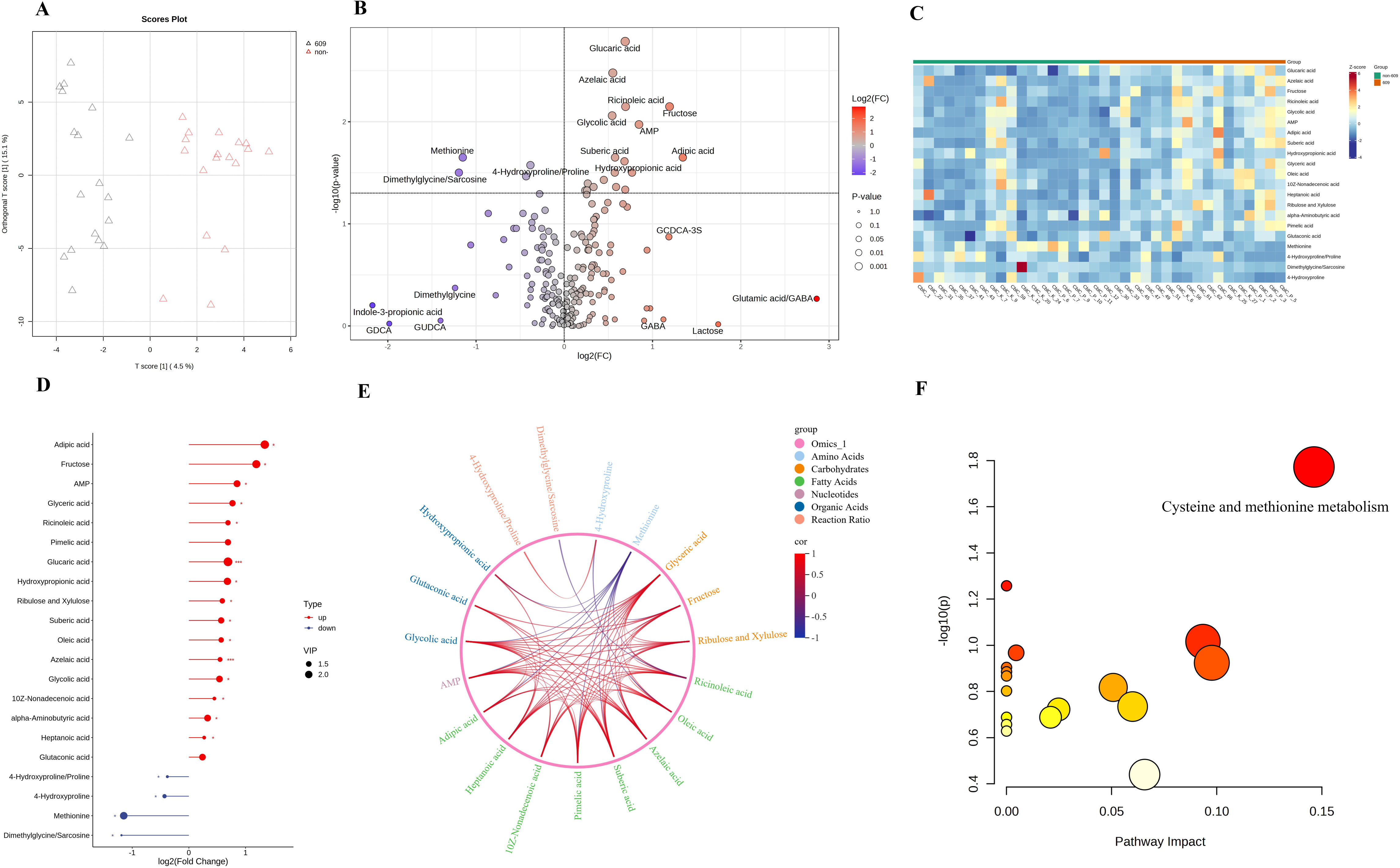
Comparative metabolome analysis of c.609G > A and non-c.609G > A carriers of cblC. (A) OPLS-DA score plot. (B) Volcano plot showing the differential metabolites. The horizontal dashed line corresponds to the significance threshold of *p* = 0.05. (C) Heatmap analysis of the potential biomarkers. Potential biomarkers selected based on OPLS-DA model and univariate statistics. Red represents upregulated metabolites, and blue represents downregulated metabolites. (D) Stickman plot showing the FC and VIP values of the potential biomarkers. (E) Correlation analysis of the potential biomarkers with significant differences (*p* < 0.05). The Chord diagram was used to represent the correlation between various biomarkers. (F) Pathway analysis based on the potential biomarkers. The x-axis represents pathway impact generated from topological analysis; the y-axis represents the corresponding p values. Con, control; cblC, patients with cobalamin C defect; FC, fold change; OPLS-DA, orthogonal partial least squares discriminant analysis; VIP, variable importance in projection.

**Fig. 6:**
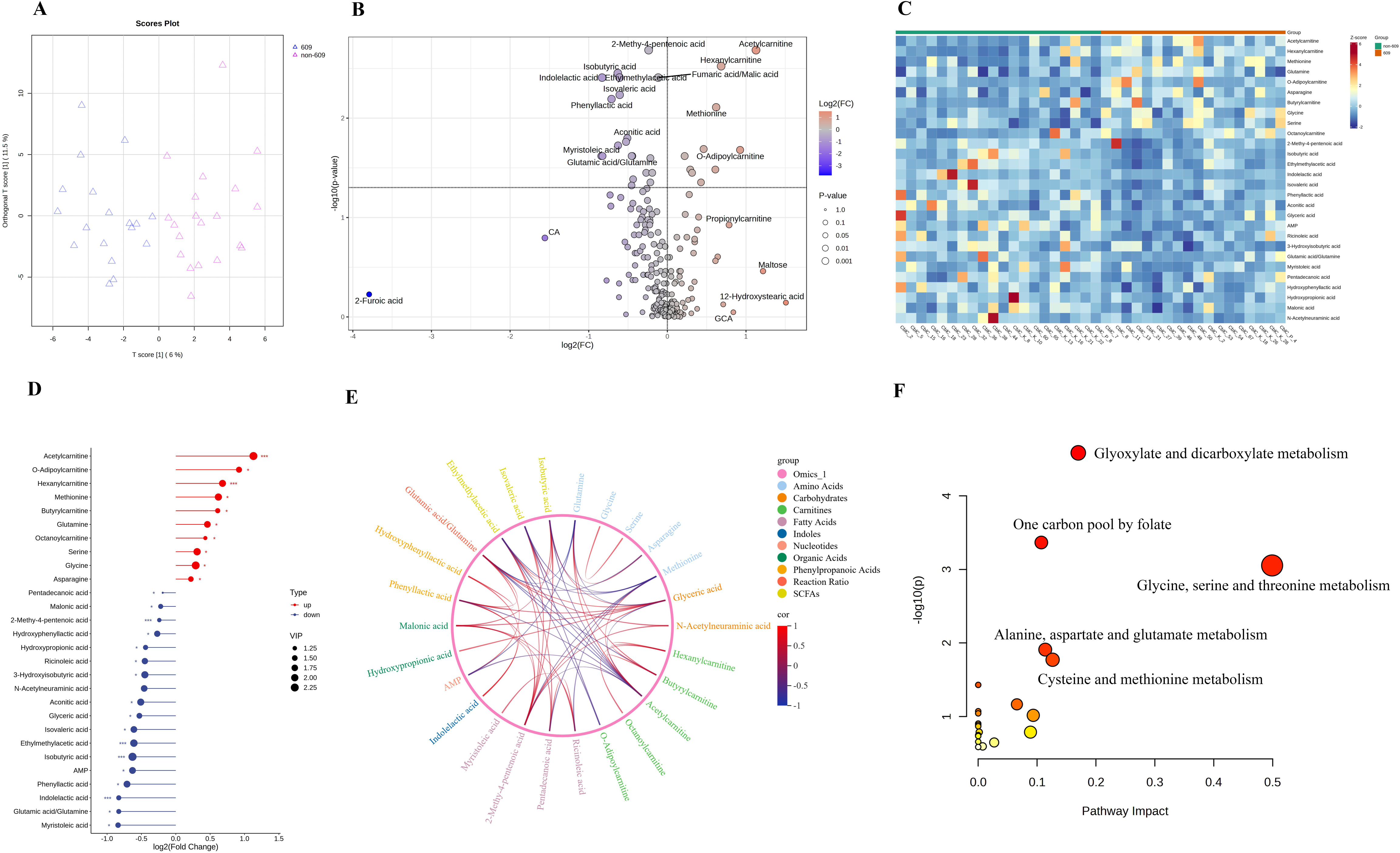
Comparative metabolome analysis of c.609G > A and non-c.609G > A carriers of cblC after treatment. (A) OPLS-DA score plot. (B) Volcano plot showing the differential metabolites. The horizontal dashed line corresponds to the significance threshold of *p* = 0.05. (C) Heatmap analysis of the potential biomarkers. Potential biomarkers selected based on OPLS-DA model and univariate statistics. Red represents upregulated metabolites, and blue represents downregulated metabolites. (D) Stickman plot showing the FC and VIP values of the potential biomarkers. (E) Correlation analysis of the potential biomarkers with significant differences (*p* < 0.05). The Chord diagram was used to represent the correlation between various biomarkers. (F) Pathway analysis based on the potential biomarkers. The x-axis represents pathway impact generated from topological analysis; the y-axis represents the corresponding p values. Con, control; cblC, patients with cobalamin C defect; FC, fold change; OPLS-DA, orthogonal partial least squares discriminant analysis; VIP, variable importance in projection.

**Fig. 7:**
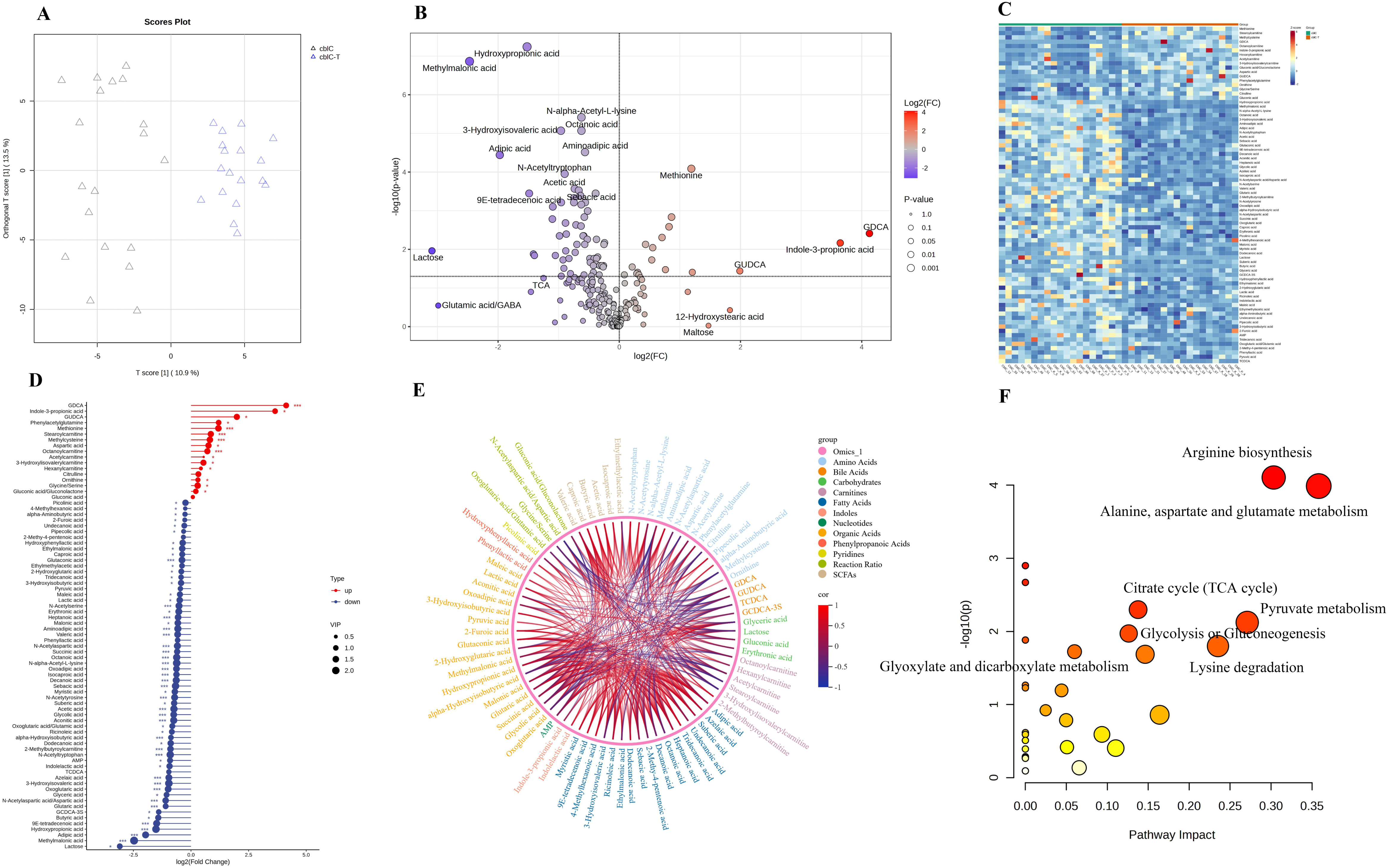
Metabolome alterations in c.609G > A carriers of cblC after treatment. (A) OPLS-DA score plot. (B) Volcano plot showing the differential metabolites. The horizontal dashed line corresponds to the significance threshold of *p* = 0.05. (C) Heatmap analysis of the potential biomarkers. Potential biomarkers selected based on OPLS-DA model and univariate statistics. Red represents upregulated metabolites, and blue represents downregulated metabolites. (D) Stickman plot showing the FC and VIP values of the potential biomarkers. (E) Correlation analysis of the potential biomarkers with significant differences (*p* < 0.05). The Chord diagram was used to represent the correlation between various biomarkers. (F) Pathway analysis based on the potential biomarkers. The x-axis represents pathway impact generated from topological analysis; the y-axis represents the corresponding p values. Con, control; cblC, patients with cobalamin C defect; FC, fold change; OPLS-DA, orthogonal partial least squares discriminant analysis; VIP, variable importance in projection.

**Fig. 8:**
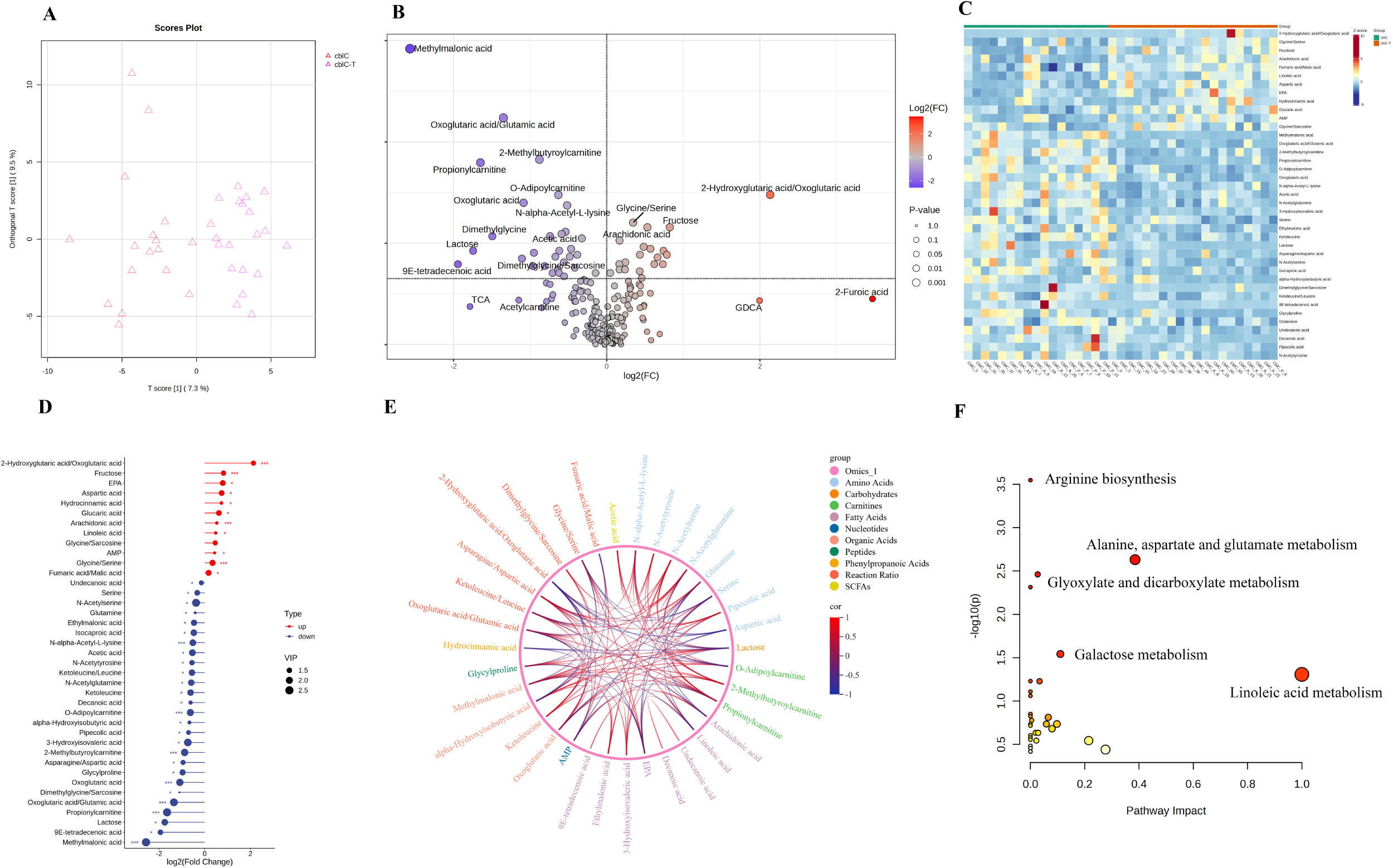
Metabolome alterations in non-c.609G > A carriers of cblC after treatment. (A) OPLS-DA score plot. (B) Volcano plot showing the differential metabolites. The horizontal dashed line corresponds to the significance threshold of *p* = 0.05. (C) Heatmap analysis of the potential biomarkers. Potential biomarkers selected based on OPLS-DA model and univariate statistics. Red represents upregulated metabolites, and blue represents downregulated metabolites. (D) Stickman plot showing the FC and VIP values of the potential biomarkers. (E) Correlation analysis of the potential biomarkers with significant differences (*p* < 0.05). The Chord diagram was used to represent the correlation between various biomarkers. (F) Pathway analysis based on the potential biomarkers. The x-axis represents pathway impact generated from topological analysis; the y-axis represents the corresponding *p* values. Con, control; cblC, patients with cobalamin C defect; FC, fold change; OPLS-DA, orthogonal partial least squares discriminant analysis; VIP, variable importance in projection.

Differential analysis of all detected metabolites based on VIP, FC, and *p* value between between c.609G > A and non-c.609G > A carriers of cblC-MMA before treatment was performed (Fig. 5A-D). Compared to the non-c.609G > A carriers, fatty acids (adipic acid, ricinoleic acid, and pimelic acid), carbohydrates (fructose, glyceric acid, glucaric acid, ribulose and xylulose), AMP, and hydroxypropionic acid were significantly upregulated in the c.609G > A carriers. 4-Hydroxyproline/proline, 4-hydroxyproline, methionine, and dimethylglycine/sarcosine were significantly down regulated (Supplementary Table 30). To gain a deeper understanding of the obtained potential biomarker, spearman rank correlation analyse was performed (Fig. 5E). The correlations of potential biomarkers with coefficients > 0.5 (*p* < 0.05) is provided in Supplementary Table 31. Methionine was negatively associated with glyceric acid, suberic acid, pimelic acid, glycolic acid, adipic acid, AMP, hydroxypropionic acid, azelaic acid, ricinoleic acid and fructose. AMP was positively associated with glycolic acid, glyceric acid, suberic acid, pimelic acid, azelaic acid, ricinoleic acid, oleic acid, adipic acid, heptanoic acid, ribulose, and xylulose. The potential biomarkers were enriched in cysteine and methionine metabolism with significant difference (Fig. 5F, Supplementary Table 32). Metablomic alterations between c.609G > A and non-c.609G > A carriers after treatment were also explored based on the OPLS-DA model and univariate analysis (Fig. 6A-D). Carnitines (acetylcarnitine, O-adipoylcarnitine, hexanylcarnitine, and butyrylcarnitine) and methionine were significantly increased in c.609G > A carriers. SCFAs (isobutyric acid, ethylmethylacetic acid, and isovaleric acid), myristoleic acid, indolelactic acid, glutamic acid/glutamine, phenyllactic acid, and AMP were significantly increased in non-c.609G > A carriers (Supplementary Table 33). Methionine was negatively associated with fatty acids (pentadecanoic acid, ricinoleic acid, and 2-methy-4-pentenoic acid) and positively associated with acetylcarnitine. Indolelactic acid was positively associated with phenyllactic acid and hydroxyphenyllactic acid (Fig. 6E, Supplementary Table 34). The potential biomarkers were mainly enriched in the following pathways: glyoxylate and dicarboxylate metabolism, one carbon pool by folate, glycine, serine and threonine metabolism, alanine, aspartate and glutamate metabolism, and cysteine and methionine metabolism (Fig. 6F, Supplementary Table 35).

Investigation of metabolomic shifts in response to treatment in c.609G > A and non-c.609G > A carriers was also performed, respectively. DBS metabolites from c.609G > A carriers before and after treatment were separated using OPLS-DA (Fig. 7A). Based on the multi- and un-variant annalysis, 78 potential biomarkers were identified. After treatment, bile acids (glycodeoxycholic acid and glycoursodeoxycholic acid), amino acids (phenylacetylglutamine, methionine, methylcysteine, and aspartic acid), carnitines (stearoylcarnitine and octanoylcarnitine), indole-3-propionic acid were significantly up-regulated in c.609G > A carriers, whereas methylmalonic acid and lactose were significantly down-regulated (Fig. 7B-D, Supplementary Table 36). Furthermore, chord diagram illustrated the correlations between different classes of metabolites in the two groups, highlighting the internal correlations within the fatty acids and bile acids. Glycodeoxycholic acid was negatively associated with alpha-hydroxyisobutyric acid, picolinic acid, and lactose. Methionine was negatively associated with some fatty acids. Methylmalonic acid was positively associated with hydroxypropionic acid, 3-hydroxyisovaleric acid, 2-methylbutyroylcarnitine, N-alpha-acetyl-L-lysine, N-acetyltryptophan, 3-hydroxyisobutyric acid, and aminoadipic acid with significant differences (Fig.7E, Supplementary Table 37). Enrichment analysis was conducted to explore the metabolic pathway alterations in c.609G > A carriers after treatment. Arginine biosynthesis, alanine, aspartate and glutamate metabolism, citrate cycle, pyruvate metabolism, glycolysis or gluconeogenesis, glyoxylate and dicarboxylate metabolism, and lysine degradation were among the most affected pathway (Fig. 7F, Supplementary Table 38). For non-c.609G > A carriers before and after treatment, OPLS-DA revealed significant metabolic changes (Fig. 8A). Based on the VIP, FC, and *p* value, 39 potential biomarkers were screened out (Fig. 8B-D, Supplementary Table 39). Of these, 12 metabolites or reaction ratios were significantly upregulated, and 27 were significantly downregulated. Carbohydrates (fructose and glucaric acid), fatty acids (EPA, arachidonic acid, and linoleic acid), and nucleotides, AMP, were significantly increased in non-c.609G > A carriers after treatment. Whereas, methylmalonic acid, 9E-tetradecenoic acid, lactose, and propionylcarnitine were significantly decreased. Furthermore, significant differences in metabolites were subjected to subsequent bioinformatic analyses. Propionylcarnitine was negatively correlated with linoleic acid, arachidonic acid, and AMP, and positively correlated with N-acetylglutamine, glutamine, and N-alpha-acetyl-L-lysine. Methylmalonic acid was negatively correlated with linoleic acid and glycine/serine, and positively correlated with lactose and serine (Fig. 8E, Supplementary Table 40). Functional analysis showed that the potential biomarkers were mainly enriched in the following pathways: arginine biosynthesis, alanine, aspartate and glutamate metabolism, glyoxylate and dicarboxylate metabolism, galactose metabolism, and linoleic acid metabolism (Fig. 8F, Supplementary Table 41).

In summary, our findings reveal that c.609G > A carriers represent a metabolically distinct state, distinguishing them from non-c.609G > A carriers. Our results underscore the importance of considering and monitoring metabolic condition when managing cblC-MMA patients carrying different gene mutations.

## Discussion

The Chinese MMA population exhibits distinct characteristics, and the combined MMA type accounts for 60%–80% of cases in China.^7^ The most prevalent *MMACHC* variant was c.609G > A in China and the c.729_730insTT in the *MMUT* gene had the highest frequency in our previous study.^10,13,18^ Moreover, the genotype–phenotype associations have also been reported for the above variants: irreversible brain disorders and poor prognosis were more common in cblC patients carrying the c.609G > A variant.^17^ However, the mechanism whereby mut deficiency and cblC deficiency with specific mutation sites presenting different clinical symptom and prognosis is unelucidated.

Comprehending metabolic reprogramming mechanisms is essential not only for obtaining mechanistic understandings of MMA advancement but also for the creation of pertinent treatments.^1,2,32,42,43,46^ In our study, we demonstrated that significant metabolic changes occured in c.729_730insTT carriers compared with non-c.729_730insTT carriers for mut-MMA patients. Metabolic alterations were also observed between c.609G > A carriers and non-c.609G > A carriers for cblC patients. The research we conducted offers more insight into the metabolic pathways that might predict the mutation sites of MMA genes, indicating that metabolites could be utilized as a promising indicator and target for the treatment of MMA subtypes.

In the current study, the comparative metabolomic analysis between mut and cblC MMA subtypes were firstly conducted. DBS metabolites with significant differences between mut and cblC were fatty acids, short-chain fatty acids, carbohydrates, and some aromatic amino acids. Functional analysis revealed that the altered pathways mainly involved in pentose phosphate pathway, tyrosine and tryptophan biosynthesis, one carbon pool by folate, amino sugar and nucleotide sugar metabolism, and fructose and mannose metabolism. The results indicate that significant metabolomic differences exist between mut and cblC subtypes, helpful for explaining the various clinical manifestations.

Metabolomic profiles of c.729_730insTT and c.609G > A carriers of MMA patients were comprehensively studied. For c.729_730insTT and non-c.729_730insTT carriers of mut-MMA patients, tryptophan related metabolites (indolelactic acid and indole-3-propionic acid), bile acids (glycohyocholic acid, taurocholic acid, and cholic acid), and fatty acids (9E-tetradecenoic acid, myristoleic acid, and linoleic acid) have undergone significant changes. Primary bile acid biosynthesis, tyrosine and tryptophan biosynthesis, and linoleic acid metabolism were the significantly enriched pathway. For c.609G > A and non-c.609G > A carriers of cblC-MMA patients, metabolomic alterations were also investigated. Differential analysis demonstrated that carbohydrates (fructose, glyceric acid, glucaric acid, ribulose and xylulose), carnitines (stearoylcarnitine and octanoylcarnitine), fatty acids (isobutyric acid, ethylmethylacetic acid, and isovaleric acid), amino acids (phenylacetylglutamine, methionine, methylcysteine, and aspartic acid) were the mainly altered metabolites between c.609G > A and non-c.609G > A carriers. Cysteine and methionine metabolism and one carbon pool by folate, and glycolysis were the primary altered pathway. Gene mutation c.729_730insTT and c.609G > A contributes to the reversal of metabolites in mut-MMA patients. Studying metabolic alterations associated with specific gene mutation could provide insights into the mechanisms underlying different clinical symptoms and prognosis of the same IEM.

Metabolomic analysis demonstrated an alteration of amino acid metabolism in the MMA patients carrying c.729_730insTT or c.609G > A, which was indicated by a disruption in the biosynthesis pathways of tyrosine and tryptophan. Tryptophan metabolism pathway, which mainly includes three metabolic pathways, namely, the 5-hydroxytryptamine, indole, and kynurenine metabolism pathways.^47^ We found that the metabolites downstream of tryptophan, such as indolelactic acid and indole-3-propionic acid, were significantly changed, which could protect the intestinal barrier function, inhibit NF-κB inflammation signaling, reduce the levels of pro-inflammatory cytokines, and protect neurons from damage.^28,48^ Stearoylcarnitine, belonged to long-chain acylcarnitines, is produced to facilitate fatty acid oxidation of the long-chain fatty acids octadecanoic acid. Stearoylcarnitine induce mitochondrial dysfunction and impaired IFNγ and GzmB production in CD8 T cells, leading to impaired anti-tumour immunity.^49^ Obvious alterations in the carnitines were observed in c.609G > A and non-c.609G > A carriers.

Microbiota-derived metabolites could directly affect the nervous system and the inhibition of neurotransmitter synthesis, resulting in neurofunctional disruption.^50^ Short-chain fatty acids, may exert anti-inflammatory functions.^51^ Short-chain fatty acids can enter systemic circulation, affect the integrity of the blood–brain barrier, and regulate neuroinflammation and neuronal function in the central nervous system. Short-chain fatty acids that enter the central nervous system are neuroactive, and, in the brain, they can directly regulate learning, memory, behavior, and disease progression.^47^ Short-chain fatty acids may be one of the mechanisms involved in the regulation of cognitive function and neuroinflammation.^52–54^ Bile acids can cross the blood–brain barrier and regulate neurological functions through their receptors. Moreover, the altered bile acid profiles are associated with amyloid, tau, and cognitive changes in patients with AD.^51,55,56^ Significant changes have occurred in the levels of fatty acids and bile acids in MMA patients carrying c.729_730insTT and c.609G > A. Given the existence of a complex bidirectional communication pathway between the gut microbiota and the nervous system; further research is warranted to elucidate the potential biological mechanisms underlying the role of the gut microbiota in the pathogenesis of MMA.

This research deepens our understanding of metabolic remodeling in MMA patients with different genetic background and highlight the influence of gene mutation on the metabolome. Additionally, our results underscore the importance of considering and monitoring metabolic alterations when managing MMA with the specific variant site. Promoting the adoption of metabolomics, are not only beneficial for clarifying MMA pathogenesis related to the variant site, but also for exploring the innovation treatments, especially for the mut-MMA, for which no effective treatment measures are available.

Although the present study demonstrated the metabolic alteration under specific gene mutation sites, it is important to interpret the causal inference with caution. Larger-scale MMA studies leveraging the geneomics and metabolomics are needed to elucidate the causal links between altered metabolites and the gene mutation sites. Further research on the integration of metabolomics and genomics of MMA patients could help elucidate the precise pathogenesis under different genetic backgrounds and provide opportunities for early intervention and the discovery of new therapeutic targets with high sensitivity and specificity. Furthermore, the use of non-targeted metabolomics platforms capable of capturing novel metabolite features could provide more comprehensive profiling in future studies, potentially uncovering previously uncharacterized metabolites that may play crucial roles associated with the hotspot variation of MMA genes.

In this study, we implemented an integrated genetic data layers and metabolomics analysis strategy to systematically investigate the metabolic biological pathways associated with MMA gene mutations (c.729_730insTT or c.609G > A mutations). Our study highlights the potential of metabolomics for discovering specific circulating biomarkers (mainly lipids, carbohydrates and some microbia related metabolites) for MMA carrying different gene mutations. The present study also provides new insights into fundamental metabolite physiology and clinical relevance. These findings and the underlying biological mechanisms need to be replicated in larger populations. Further studies are needed to shed light on the mechanisms underlying these potential biomarkers and to further investigate their potential clinical applications.

## Contributors

H.J.Z.: Writing – review & editing, Writing–original draft, Visualization, Validation, Supervision, Software, Project administration, Methodology, Investigation, Formal analysis, Conceptualization. Y.X.D., Y.D., and T.C.: Writing–review & editing, Writin –original draft, Methodology, Data curation, Conceptualization. B.H.Z., Y.C., S.Y.C., X.Z., and F.X.: Writing–review & editing, Writing–original draft, Methodology. L.L.H., L.L.L., and W.J.Q.: Writing–review & editing. H.W.Z., and X.F.G.: Methodology, Data curation. L.S.H.: Writing–review & editing, Supervision, Resources, Investigation, Conceptualization. All authors read and approved the final manuscript.

## Data sharing statement

The raw data in this study is available from the corresponding author upon request.

## Acknowledgements

This work was supported by the Beijing Medical Reward Foundation (No. YXJL-2025-0456-0051).

## Appendix A. Supplementary data

Supplementary data to this article can be found online.

